# Assessment of long-term trends in a threatened grassland bird community using daily bird lists

**DOI:** 10.1101/2022.10.31.514489

**Authors:** Akshay Bharadwaj, Sarang Mhamane, Priti Bangal, Tarun Menon, Kavita Isvaran, Suhel Quader

**Affiliations:** Centre for Ecological Sciences, Indian Institute of Science, Bangalore 560012; Nature Conservation Foundation. 1311, “Amritha”, 12th Main. Vijayanagar 1st Stage. Mysore 570 017

**Keywords:** Long-term monitoring, Daily checklists, Grassland birds, Seasonal trends

## Abstract

Open Natural Ecosystems (ONEs), such as tropical grasslands, are among the most threatened habitats on Earth today. The long-term monitoring of ONEs is an important research domain that is essential for understanding anthropogenic impacts and facilitating conservation action. Using a simple day-listing method over a 13-year period, we studied species trends in a central Indian grassland-agriculture mosaic experiencing several land-use changes. Our results indicate that some grassland species (such as the Great Indian Bustard *Ardeotis nigriceps*) show steep declines during the study period, while other generalist species (such as the Indian Peafowl *Pavo cristatus*) show an increasing trend. Daily listing also reveals distinct seasonal patterns, and we discuss the Great Indian Bustard and Western Marsh Harrier *Circus aeruginosus* as examples. Our study highlights the utility of consistent checklist surveys to monitor population trends of bird communities within a changing landscape.

## Introduction

Grasslands are one of the most threatened ecosystems on Earth today, a condition they share with other open natural ecosystems (ONEs) (Madhusudan and Vanak 2021). Their favourable topographic features and fertile soils have made grassland habitats the most extensively modified ecosystem by humans (Henwood 1998). These modifications and the resulting fragmentation have led to increased habitat heterogeneity and can severely threaten native grassland species (Punjabi et al. 2013).

Local bird communities are good indicators of ecosystem health and functioning (Gregory and van Strien 2010). Therefore, the study of bird communities can be useful in habitat assessment and conservation planning of a region. Grassland birds are often specialised to (or have a preference for) open habitats. Many grassland specialist birds are either ground-nesting or build small nests, which are camouflaged in grasses and reeds to avoid nest predation (Fogarty et al. 2017). Specialised species, such as the Red-necked Falcon *Falco chicquera*, show a high degree of physiological and behavioural adaptation to the grassland habitat (Ali 1990). Consequently, these species are highly sensitive to habitat features (such as vegetation type), making them vulnerable to land-use changes (such as the degree of livestock grazing; Kher and Dutta 2021).

Grassland bird communities face a multitude of threats from anthropogenic change today. Despite this, there is inadequate funding, research, and conservation effort focused on grasslands (Madhusudan and Vanak 2021). Although many grassland birds are known to be negatively affected by disturbances, low-intensity agro-pastoral lands can, in some cases, supplement protected areas in conserving grassland species and bird communities (Dutta and Jhala 2014; Kher and Dutta 2021).

Here, we use a simple checklist method to study long-term trends in bird communities within the grasslands of Nannaj, Maharashtra, India — a region with a complex interplay between biodiversity conservation and economic interests (Narwade and Rahmani 2020). Our long-term study reveals several interesting trends in reporting rates of the regional bird community. We also highlight differences in local species trends at Nannaj with their national trends. Lastly, we explore the real-world utility of a simple checklist methodology performed by an individual, a committed observer consistently over a long time.

### Study area and its current land use distribution

We documented birds within a study area bounded by the five villages of Vadala, Akolekati, Karamba, Mardi and Narotewadi, and centred around Nannaj village (17.836 **°**N, 75.851 **°**E) in the Solapur district of Maharashtra. The annual precipitation in the region is less than 750 mm, and the semi-arid climate has a distinct seasonality; long, intense summers (March-June) are followed by a rainy season (monsoon; July-October), after which the winter season (November-February) leads back into the summer (Krishna et al. 2016).

The study area, much like the larger landscape, is an evolving mosaic of protected native grasslands, afforested woodland plots, communal and private grazing lands, urban settlements, and agricultural land (Punjabi et al. 2013; Krishna et al. 2016; Narwade and Rahmani 2020). Among the main crops grown in the region are jowar (millets) and groundnut. Most farmers also own cattle, which are often allowed to graze freely in the grasslands. Our study area encompasses parts of the Great Indian Bustard Sanctuary, a protected area created to conserve the region’s native grasslands and, especially, the critically endangered Great Indian Bustard (*Ardeotis nigriceps*), locally known as *Maldhok*.

## Methods

All field data from 2009 to 2021 were collected by SM, a seasoned birdwatcher who is familiar with all the bird species in his landscape. SM maintained a daily bird attendance register containing commonly seen and easily identifiable species found in the study area (**Table 2**). There are 199 bird species recorded in Nannaj on the eBird Database (Sullivan et al. 2009; eBird 2021). Of these, we began by monitoring 40 bird species in 2009, and added a further 7 species in 2013. For each of the species on the master list, SM used a physical register to mark those seen through the course of a day, while going about his usual fieldwork routine, with no fixed route being followed. The study area depicted in Figure 1 shows the region SM typically covered as part of his daily activities. Only birds observed within the study area were included in the dataset.

**Figure 1:**
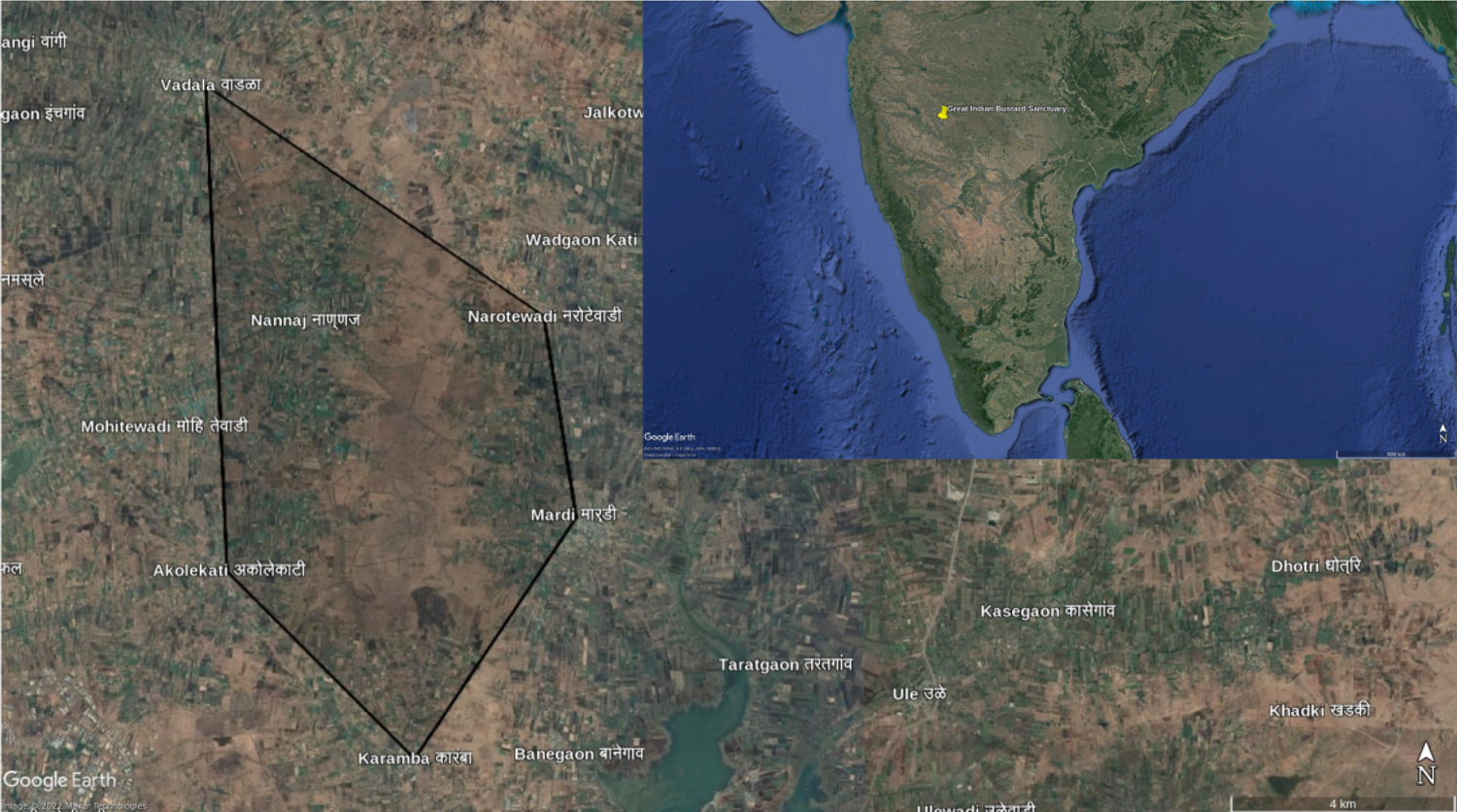
(*inset*) The location of the study area within a larger map of the Indian subcontinent; (*main*) The study area (shaded darker) is roughly a polygon with its vertices being at adjoining villages. Lighter shades of brown (in the centre) show grassland habitats within the Great Indian Bustard Sanctuary, while the darker and greener patches show agricultural lands.

**Figure 2:**
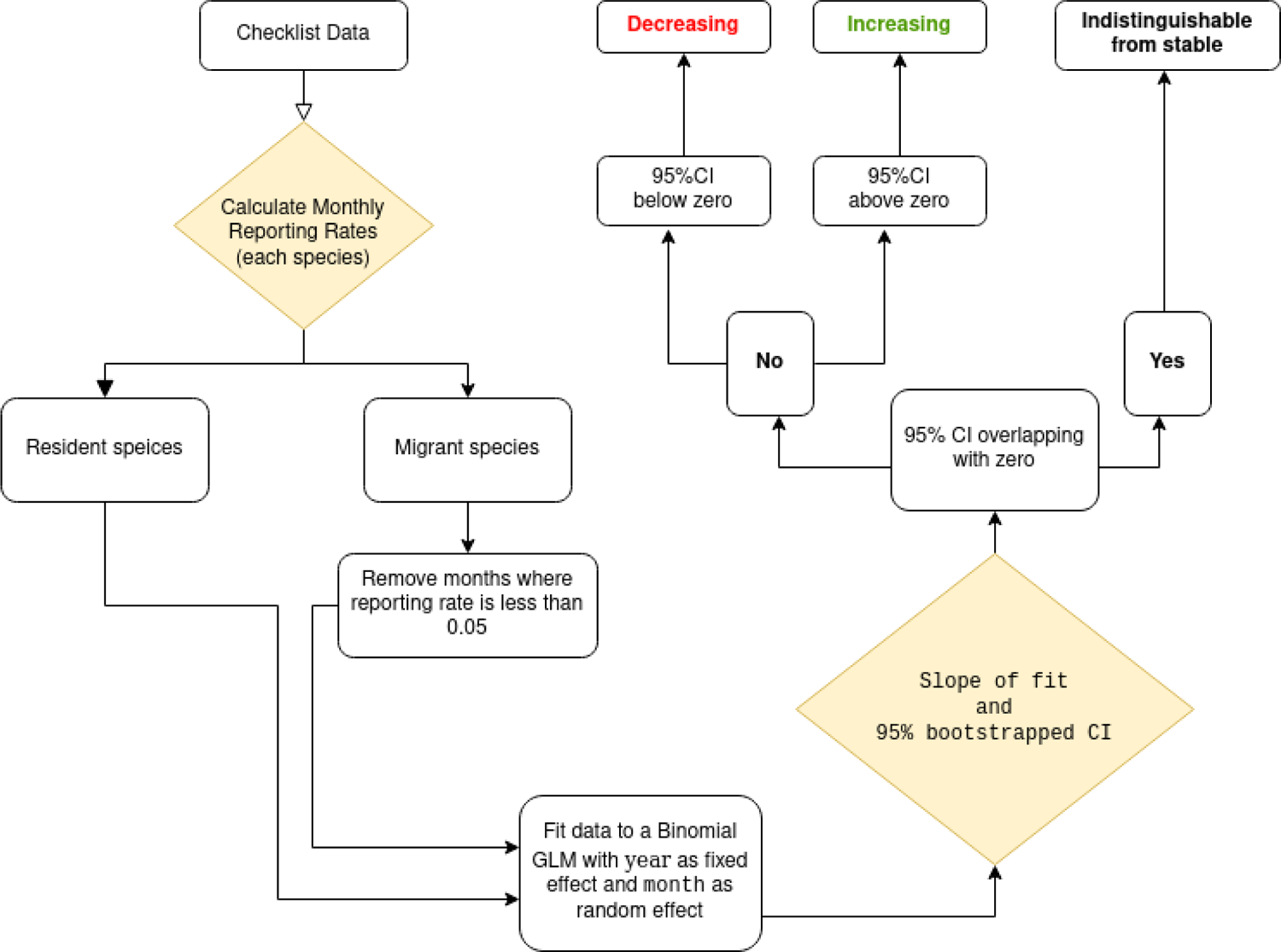
A schematic representation of the analysis pipeline.

The bird attendance register was filled at the end of each day by putting a tickmark against all species that were seen or heard during the course of that day. Species that were not seen or heard were marked with an ‘X’ mark, representing a non-detection. If the identification was uncertain, the species was recorded as undetected for the day. This routine was repeated whenever SM conducted fieldwork within the area; therefore, it mostly excluded Sundays and public holidays.

### Analysis

To analyse the collected data, the handwritten registers were digitised into a spreadsheet (row-column) format. Subsequently, the data were analysed using the following packages in R (R Development Core Team 2013) — dplyR for data handling (Wickham et al. 2019); lubridate for handling dates (Grolemund and Wickham 2011); and ggplot2 for data visualisation (Wickham 2016).

We did not have information on the number of individuals that were seen by SM. Rather, we had data on presence and absence (more accurately, detection and non-detection). Assuming that the probability of detection rises asymptotically with population density (Altwegg and Nichols 2019), we calculated and used the *reporting rate* (the fraction of checklists containing a species in a given time period) as a relative index of population density over time. Importantly, we can do this because we do not compare absolute reporting rates between species. Rather, we are only interested in studying how the reporting rates of each species change over time, in other words, we examine the trends in reporting rate for individual species. Further, because of the asymptotic nature of the relationship between reporting rate and absolute population density, reporting rate is expected to be a particularly sensitive index at low and medium densities; at high densities, reporting rate is likely to underestimate underlying population change.

For resident species, we calculated monthly reporting rates, i.e., the fraction of days in the month when a particular species was observed. Winter migrants are absent during the summer at Nannaj, and therefore for these species, we used only those months in each year with reporting rate above 0.05, thereby excluding months where the species was largely or completely absent.

The Black Redstart *Phoenicurus ochruros* was recorded only sporadically and hence removed from any further analyses. For visualisation purposes alone (in Figure 4), we calculated the annual reporting rates, which were the average of all monthly reporting rates for the species in each year.

To quantify and examine trends over years, we used the lme4 package (Bates et al. 2015) in R to fit a binomial generalised linear mixed model for each species with the month of the year as the random effect. The reason we did this was that the same month (for instance, January) is expected to have similar characteristics in terms of species occurrence across years in comparison with another month (for instance, May). A key assumption we make is that while the detection probability of a species may vary between months, it does not change across the years of the study period. In brief, this method calculates an average month-specific trend for a species across years, taking into account that different months might have different baseline reporting rates.

The slope estimates from the GLMMs for each species were tabulated along with 95% confidence intervals, calculated through robust non-parametric bootstrapping within months across years (i.e, with replacement). This allowed us to visualise broad trends (increasing, indistinguishable from stable, or declining; examples shown in Figure 4) in the reporting rate and a confidence interval around a slope estimate (shown in Figure 5). Any species whose CI overlapped zero was categorised as *Indistinguishable from stable*, while those with CIs fully above and fully below zero were categorised as *Increasing* and *Declining,* respectively. The model fitted to each species was as follows:

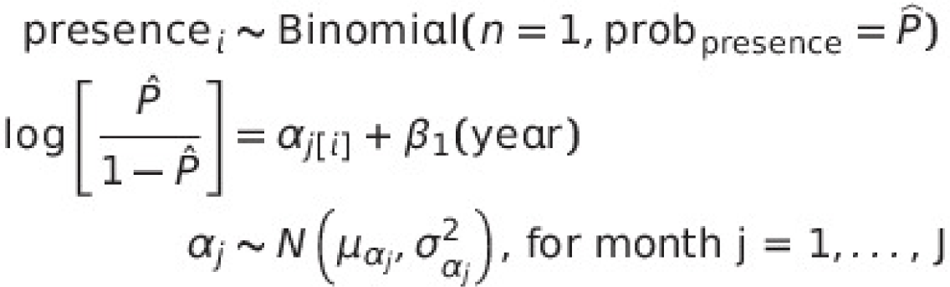

In addition to examining annual trends, we also investigated seasonal changes in bird reporting rates for select species by calculating reporting rates for each month, averaged across years. 95% confidence intervals for the month estimates were calculated using the Agresti-Coull method (Brown et al. 2001).

To understand how trends might differ among different kinds of species, we also classified species into different habitat specialisation guilds. This classification was based largely on the *State of India’s Birds* report (SoIB 2020), supplemented with information from *The Book of Indian Birds* (Ali 1990) and the *Birds of the World* database (Billerman et al. 2022). Definitions for each habitat guild are in Table 1. To visualise temporal changes in reporting rates for a guild as a whole (Figure 3), we first calculated the reporting rate of each species in each year (as shown above). For each year, we then calculated the mean reporting rates and 95% CIs across all the species in that guild.

**Figure 3:**
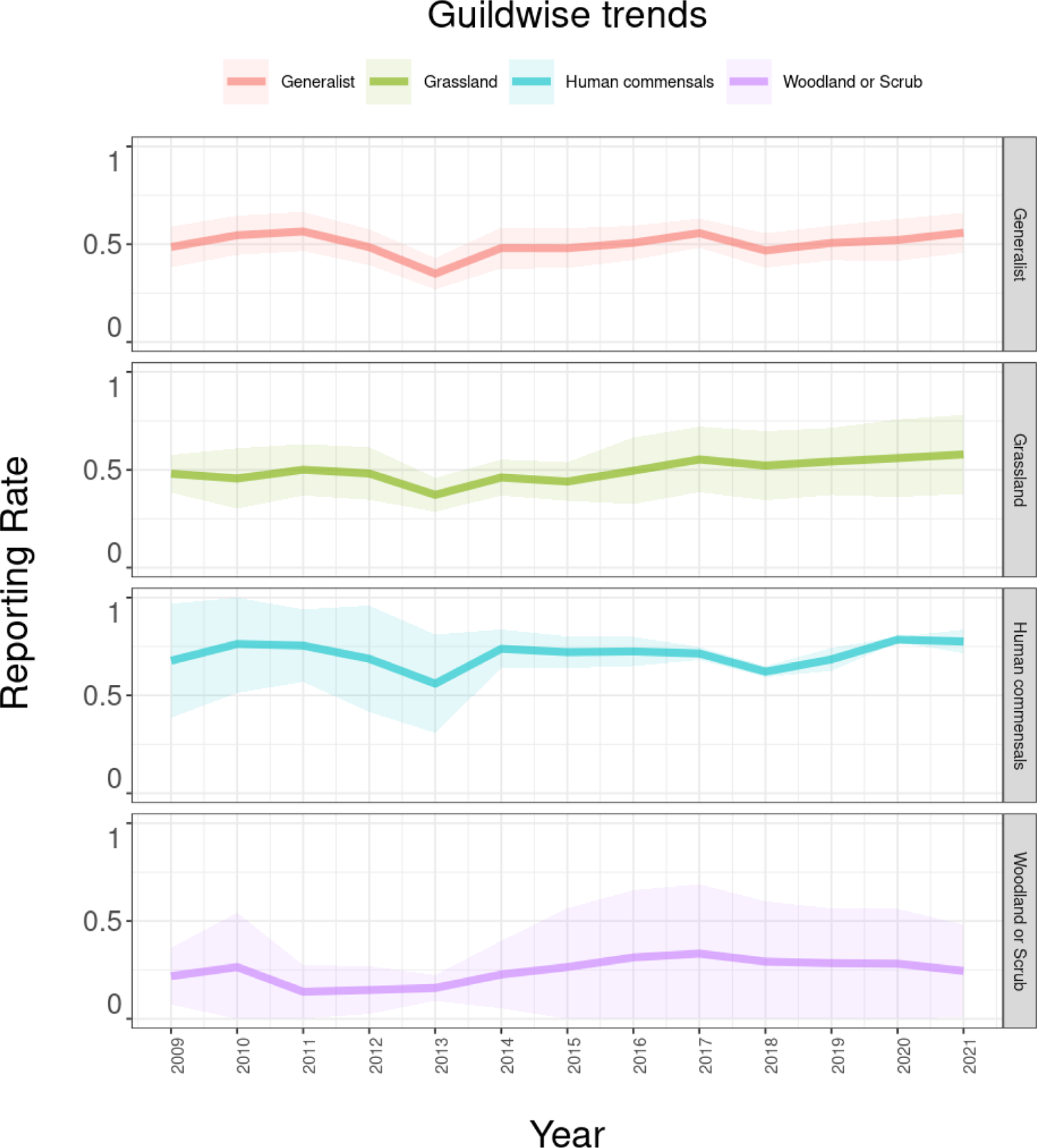
Guild-wise trends in reporting rates across the study period (2009-2020). The shaded region represents the 95% confidence interval around the mean.

**Table 1:**
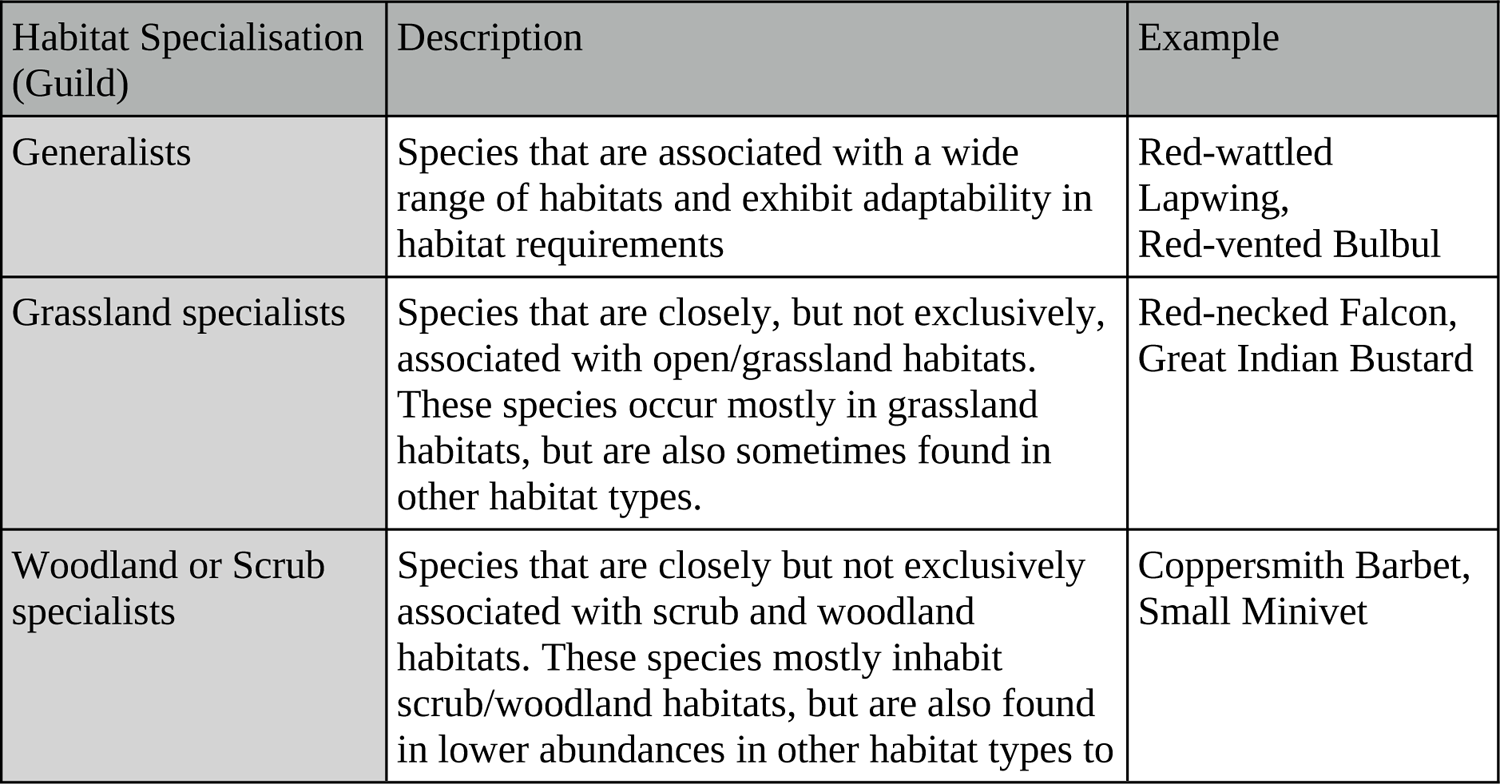
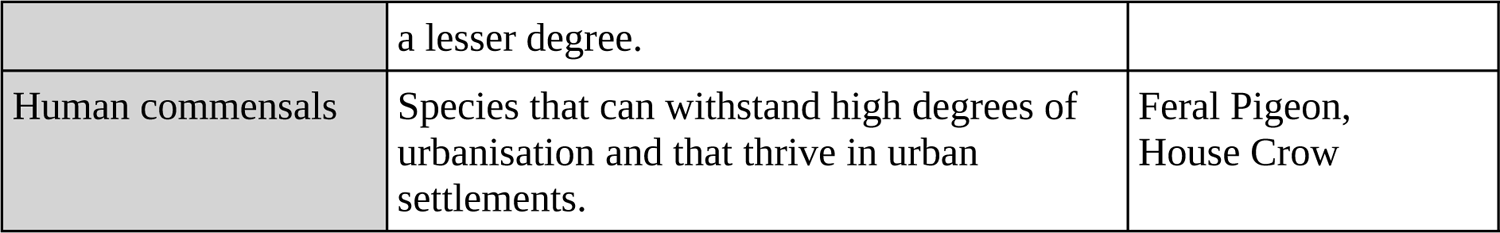
Description and example species of each guild in the study

| Habitat Specialisation (Guild) | Description | Example |
| --- | --- | --- |
| Generalists | Species that are associated with a wide range of habitats and exhibit adaptability in habitat requirements | Red-wattled Lapwing,<br>Red-vented Bulbul |
| Grassland specialists | Species that are closely, but not exclusively, associated with open/grassland habitats. These species occur mostly in grassland habitats, but are also sometimes found in other habitat types. | Red-necked Falcon,<br>Great Indian Bustard |
| Woodland or Scrub specialists | Species that are closely but not exclusively associated with scrub and woodland habitats. These species mostly inhabit scrub/woodland habitats, but are also found in lower abundances in other habitat types to | Coppersmith Barbet,<br>Small Minivet |
|  | a lesser degree. |  |
| Human commensals | Species that can withstand high degrees of urbanisation and that thrive in urban settlements. | Feral Pigeon,<br>House Crow |

**Table 2:**
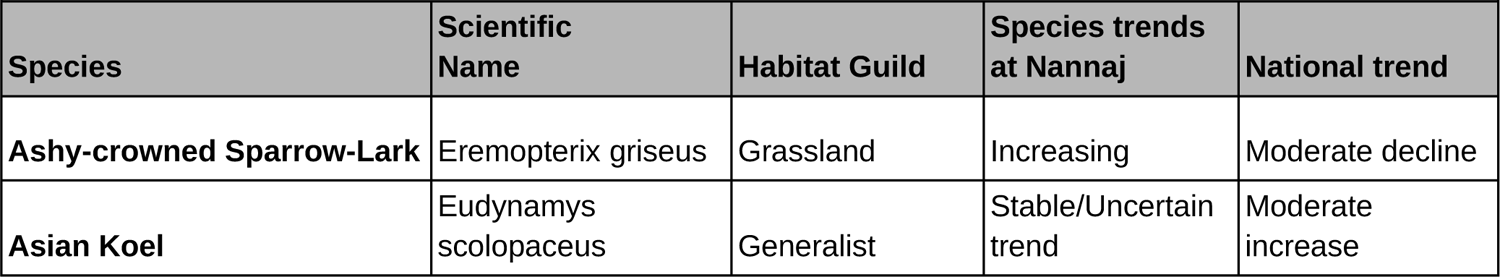
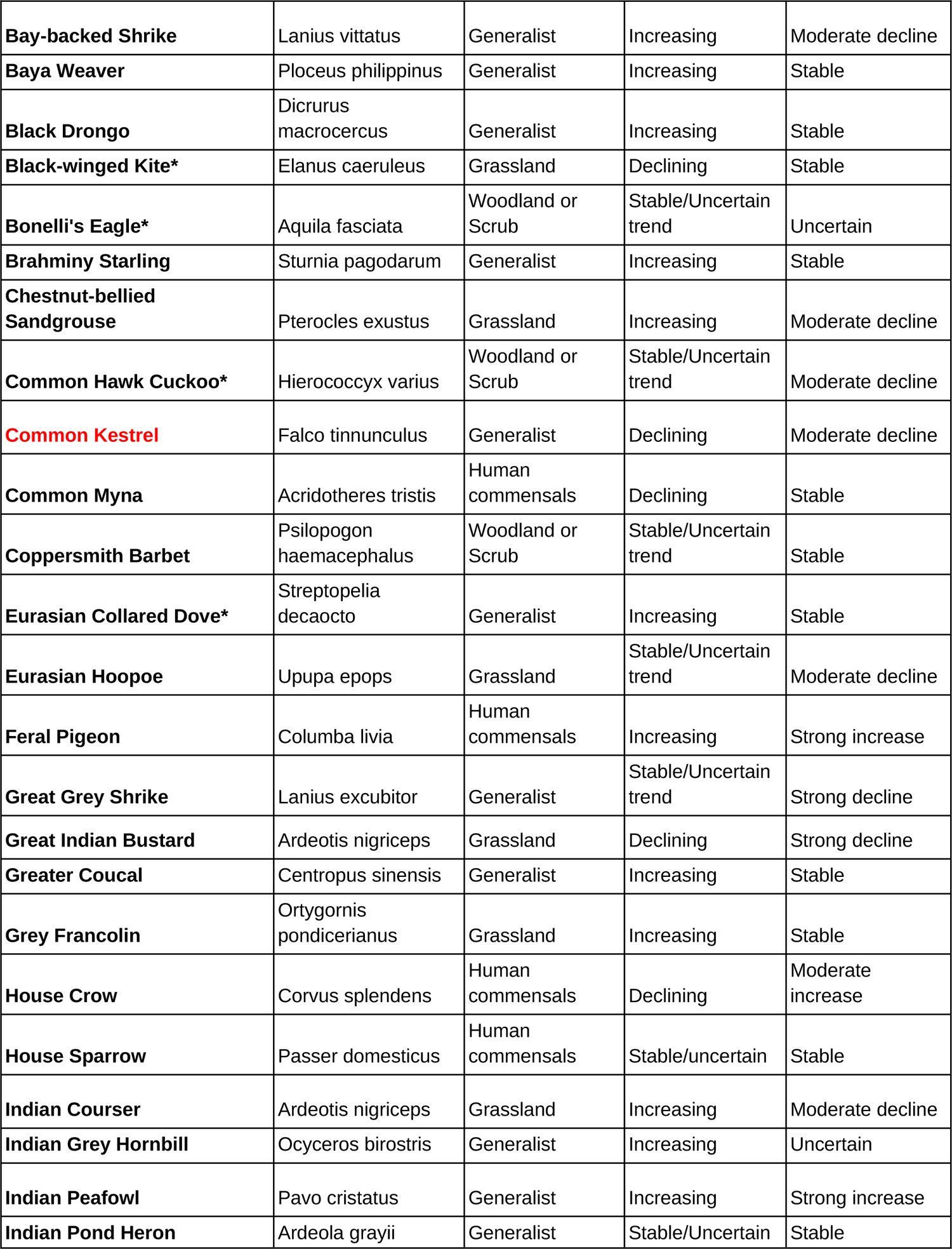
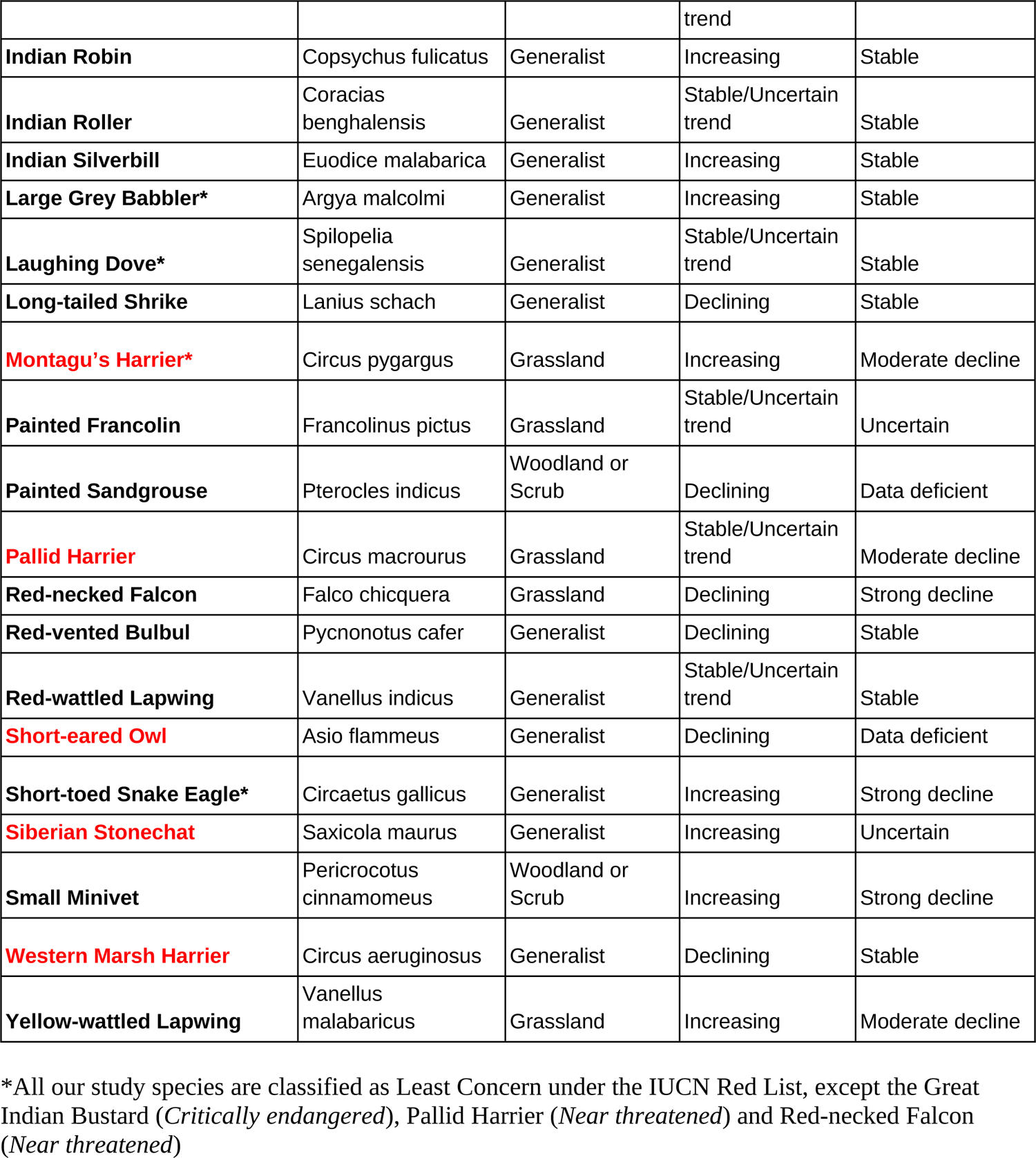
Species-level changes in reporting rates over time, habitat guilds, local species trends and national species trends (SoIB 2020) of commonly seen bird species at Nannaj. Species in red are winter migrants at Nannaj, and those with an asterisk (*) have been monitored only since 2013.

| Species | Scientific Name | Habitat Guild | Species trends at Nannaj | National trend |
| --- | --- | --- | --- | --- |
| Ashy-crowned Sparrow-Lark | Eremopterix griseus | Grassland | Increasing | Moderate decline |
| Asian Koel | Eudynamys scolopaceus | Generalist | Stable/Uncertain trend | Moderate increase |
| **Bay-backed Shrike** | *Lanius vittatus* | Generalist | Increasing | Moderate decline |
| **Baya Weaver** | *Ploceus philippinus* | Generalist | Increasing | Stable |
| **Black Drongo** | *Dicrurus macrocercus* | Generalist | Increasing | Stable |
| **Black-winged Kite\*** | *Elanus caeruleus* | Grassland | Declining | Stable |
| **Bonelli's Eagle\*** | *Aquila fasciata* | Woodland or Scrub | Stable/Uncertain trend | Uncertain |
| **Brahminy Starling** | *Sturnia pagodarum* | Generalist | Increasing | Stable |
| **Chestnut-bellied Sandgrouse** | *Pterocles exustus* | Grassland | Increasing | Moderate decline |
| **Common Hawk Cuckoo\*** | *Hierococcyx varius* | Woodland or Scrub | Stable/Uncertain trend | Moderate decline |
| **Common Kestrel** | *Falco tinnunculus* | Generalist | Declining | Moderate decline |
| **Common Myna** | *Acridotheres tristis* | Human commensals | Declining | Stable |
| **Coppersmith Barbet** | *Psilopogon haemacephalus* | Woodland or Scrub | Stable/Uncertain trend | Stable |
| **Eurasian Collared Dove\*** | *Streptopelia decaocto* | Generalist | Increasing | Stable |
| **Eurasian Hoopoe** | *Upupa epops* | Grassland | Stable/Uncertain trend | Moderate decline |
| **Feral Pigeon** | *Columba livia* | Human commensals | Increasing | Strong increase |
| **Great Grey Shrike** | *Lanius excubitor* | Generalist | Stable/Uncertain trend | Strong decline |
| **Great Indian Bustard** | *Ardeotis nigriceps* | Grassland | Declining | Strong decline |
| **Greater Coucal** | *Centropus sinensis* | Generalist | Increasing | Stable |
| **Grey Francolin** | *Ortygornis pondicerianus* | Grassland | Increasing | Stable |
| **House Crow** | *Corvus splendens* | Human commensals | Declining | Moderate increase |
| **House Sparrow** | *Passer domesticus* | Human commensals | Stable/uncertain | Stable |
| **Indian Courser** | *Ardeotis nigriceps* | Grassland | Increasing | Moderate decline |
| **Indian Grey Hornbill** | *Ocyrceros birostris* | Generalist | Increasing | Uncertain |
| **Indian Peafowl** | *Pavo cristatus* | Generalist | Increasing | Strong increase |
| **Indian Pond Heron** | *Ardeola grayii* | Generalist | Stable/Uncertain | Stable |
|  |  |  | trend |  |
| **Indian Robin** | *Copsychus fulicatus* | Generalist | Increasing | Stable |
| **Indian Roller** | *Coracias benghalensis* | Generalist | Stable/Uncertain trend | Stable |
| **Indian Silverbill** | *Euodice malabarica* | Generalist | Increasing | Stable |
| **Large Grey Babbler\*** | *Argya malcolmi* | Generalist | Increasing | Stable |
| **Laughing Dove\*** | *Spilopelia senegalensis* | Generalist | Stable/Uncertain trend | Stable |
| **Long-tailed Shrike** | *Lanius schach* | Generalist | Declining | Stable |
| **Montagu's Harrier\*** | *Circus pygargus* | Grassland | Increasing | Moderate decline |
| **Painted Francolin** | *Francolinus pictus* | Grassland | Stable/Uncertain trend | Uncertain |
| **Painted Sandgrouse** | *Pterocles indicus* | Woodland or Scrub | Declining | Data deficient |
| **Pallid Harrier** | *Circus macrourus* | Grassland | Stable/Uncertain trend | Moderate decline |
| **Red-necked Falcon** | *Falco chicquera* | Grassland | Declining | Strong decline |
| **Red-vented Bulbul** | *Pycnonotus cafer* | Generalist | Declining | Stable |
| **Red-wattled Lapwing** | *Vanellus indicus* | Generalist | Stable/Uncertain trend | Stable |
| **Short-eared Owl** | *Asio flammeus* | Generalist | Declining | Data deficient |
| **Short-toed Snake Eagle\*** | *Circaetus gallicus* | Generalist | Increasing | Strong decline |
| **Siberian Stonechat** | *Saxicola maurus* | Generalist | Increasing | Uncertain |
| **Small Minivet** | *Pericrocotus cinnamomeus* | Woodland or Scrub | Increasing | Strong decline |
| **Western Marsh Harrier** | *Circus aeruginosus* | Generalist | Declining | Stable |
| **Yellow-wattled Lapwing** | *Vanellus malabaricus* | Grassland | Increasing | Moderate decline |

\*All our study species are classified as Least Concern under the IUCN Red List, except the Great Indian Bustard (*Critically endangered*), Pallid Harrier (*Near threatened*) and Red-necked Falcon (*Near threatened*)

With this information in hand, we asked three broad questions:

1. How did the reporting rates of habitat guilds change over the study period?
2. How did the species-level reporting rates change over time?
3. Within a single species, can we observe seasonal changes/patterns in reporting rates?

## Results

We monitored 45 bird species, including seven migratory species, in 4324 days of survey over 13 years. The slope estimates and bootstrapped confidence intervals of the trends of each species (examples shown in Figure 4) across the study period are shown in Figure 5, where the species are ordered by their overall mean reporting rates.

**Figure 4:**
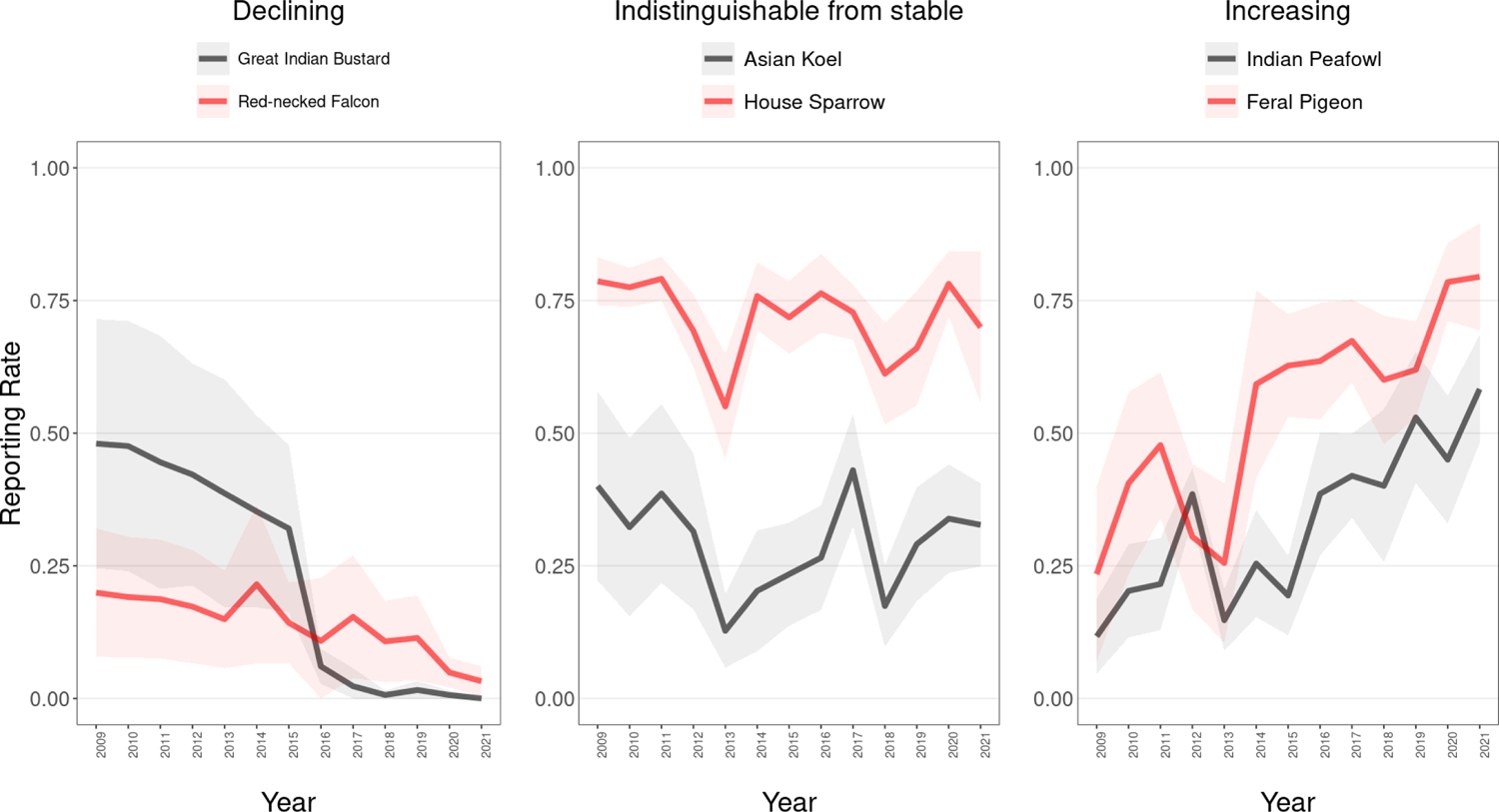
Examples of different species showing declining, indistinguishable from stable and increasing trends in their annual reporting rates over time. The shaded regions represent 95% confidence interval around the mean.

Table 2 shows a summary of species-wise classifications and results. All species common names used are from the *India list* published by *Indian Birds* (Praveen et al. 2016).

Guild-wise trends across the study period indicate that reporting rates of various habitat guilds in the region have been largely stable (**Figure 3**).

Among grassland species, smaller-bodied and more diet-generalist species (such as the Ashy-crowned Sparrow Lark *Eremopterix griseus;* **Figure 5**) showed minor increases in reporting rates. Meanwhile, larger-bodied specialist species (such as the Great Indian Bustard and Red-necked Falcon) showed strong, consistent declines throughout the study period (**Figure 5**).

**Figure 5:**
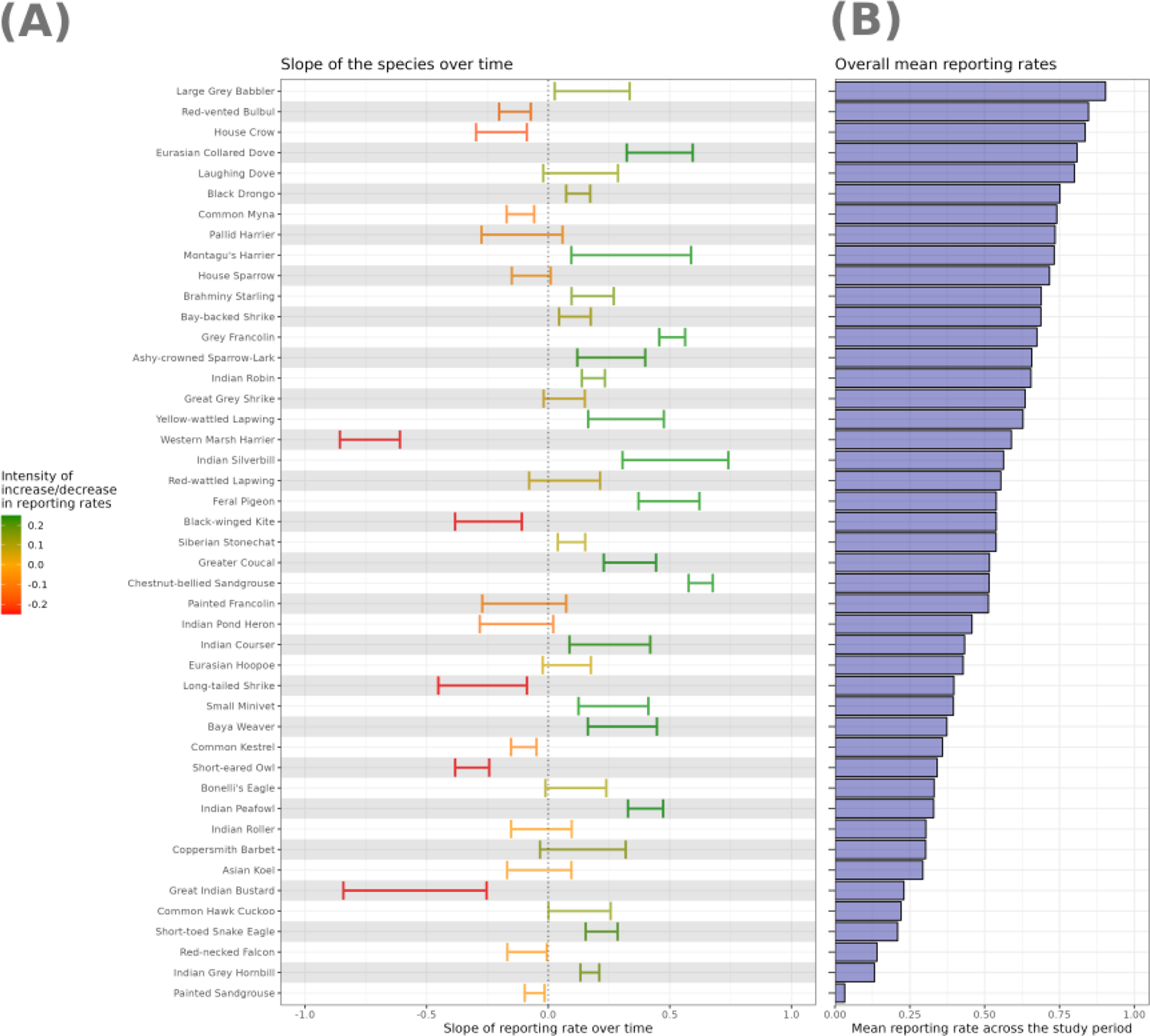
(A) Estimates of the slopes of reporting rate over time (across the study period), as derived from GLMM analysis. Error bars are 95% CIs. Colours reflect the magnitude of the estimated slope, as described in the accompanying key. (B) Overall mean reporting rate of each species across all years.

Generalist species, on the whole, had indistinguishable from stable or increasing trends over time (**Figure 5**). However, some species with a preference for more open habitats (such as the Western Marsh Harrier; Kitowski 2007) showed considerable declines over time.

Woodland/scrub species (such as the Small Minivet *Pericrocotus cinnamomeus*) showed increasing reporting rates over time, except the Painted Sandgrouse *Pterocles indicus*, a relatively rare species of scrub habitat, which appears to have experienced a considerable decline (**Figure 5**).

Interestingly, some human commensals, which often inhabit urban and human-modified landscapes (such as the Common Myna and House Crow), have witnessed moderate declines in reporting rates, while another commensal, the Feral Pigeon, has increased considerably (**Figure 5**).

### Studying seasonal trends in bird populations

To investigate what can be learnt from this dataset about seasonal trends using our methodology, we examine two species - (a) Western Marsh Harrier, a migratory raptor that travels over long distances between central Asia and the Indian subcontinent; and (b) the Great Indian Bustard, a grassland-specialist and flagship species that shows seasonal local movements.

India is one of the largest wintering grounds for harriers in the world. As the only raptors in the world that nest on the ground, harriers are adapted to living in open landscapes such as grassland-marsh mosaics. Furthermore, as one of the top predators in the grassland food chain, Harriers can serve as an indicator of ecosystem health (Verma 1996).

The Western Marsh Harrier is a common, widespread winter visitor to India. It is often found in a mosaic of marshland, agricultural fields and grassy plains. Throughout our study, we see a steady decline in winter reporting rates of this species (Figure 6). This trend is consistent with other long-term studies of roost counts of harriers in India (Ganesh and Prashanth 2018).

**Figure 6:**
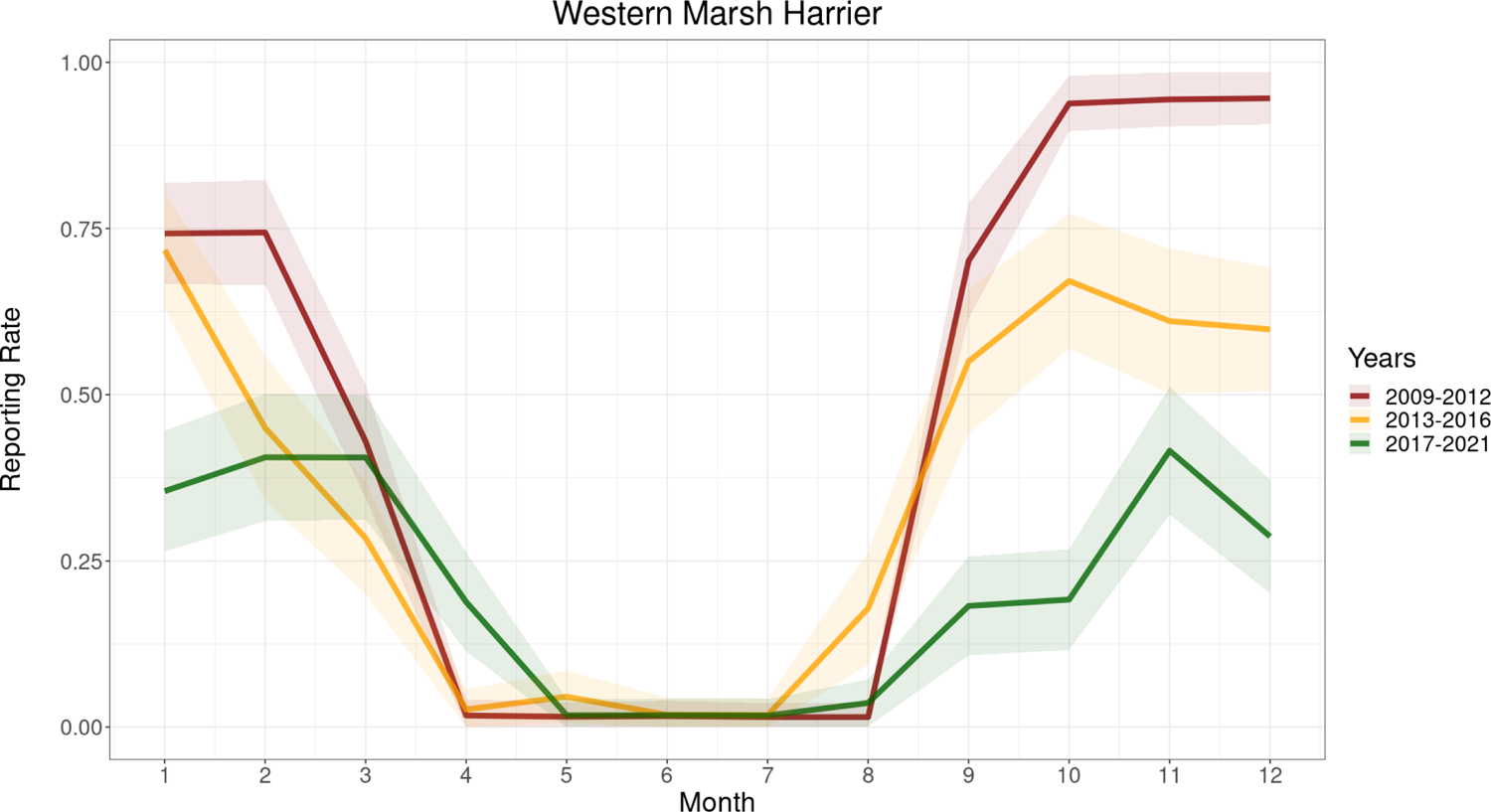
Monthly trends in reporting rates of the Western Marsh Harrier throughout the study period, with every four years grouped together for clearer visualisation. This is a species with a preference for open habitats. There is an evident decline in winter reporting rates in each consecutive four-year period.

In the early years of this study, the Great Indian Bustard showed a distinct seasonality, with reporting rates being highest from July to October. Overall reporting rates have declined steadily, and in recent years the reporting rate is so low that no seasonality is apparent any more (Figure 7).

**Figure 7:**
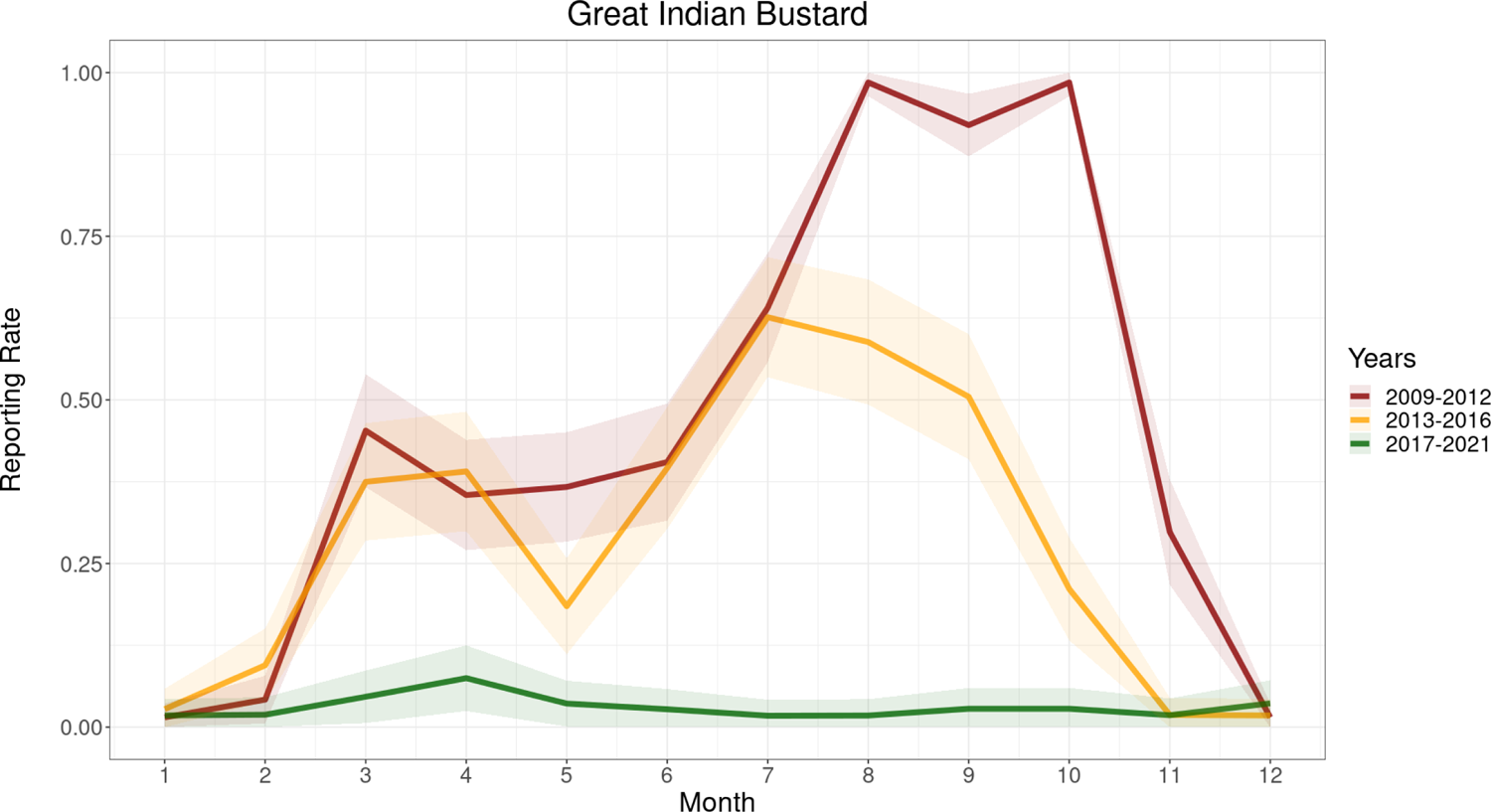
Monthly trends in reporting rates of the Great Indian Bustard throughout the study period, with every four years grouped together for clarity in visualisation. There is a clear seasonality in the bustard activity within the study area. In recent years, reports of the species have fallen to near zero.

## Discussion

Our study used a simple checklist-based method to monitor species trends over 13 years. This method can be implemented by anyone with a basic knowledge of bird identification without the need for intensive training in more sophisticated protocols. While quick and easy, such a method is not suited for monitoring population densities taking into account detectability. In other words, we cannot compare the reporting rates across different species (because they will often differ in detectability), rather we focus on looking at within-species trends. For inter-species comparisons, more detailed protocols like distance-sampling-based transects or point counts are needed. Its simplicity, however, makes the checklist method a widely implementable procedure to examine general trends.

Despite changes in land use within the study area, including increasing urbanisation and extensive agricultural expansion in the region (Narwade and Rahmani 2020), many species show no discernible increases or decreases in reporting rates over the study period. This suggests that the region continues to support a large fraction of its common species.

A handful of sensitive species, however, mostly large-bodied grassland specialists, show steady declines (Figure 5), possibly due to the loss of native grassland habitat (Madhusudan and Vanak 2021). Among human commensal species, Feral Pigeon shows a tremendous increase, while Common Myna and House Crow show a decreasing local trend (Figure 5). Below, we take a closer look at trends in selected groups and species.

### A. Grassland species

Grassland species varied in their trends over time. Small-bodied grassland generalists such as the Ashy-crowned Sparrow Lark showed an increase in reporting rates over time (Figure 5). The Yellow-wattled Lapwing *Vanellus malabaricus*, a species with a high affinity for grassland habitats (Sethi et al. 2010), also showed a moderate increase in reporting rate (Figure 5). This is in contrast to its steadily declining national population trends (SoIB 2020), and indicates the potential of the Nannaj grasslands in conserving this increasingly threatened species. Similarly, the Indian Courser *Cursorius coromandelicus* showed increases in their reporting rates (Figure 5), contrasting with their strong declines at the national scale (SoIB 2020). Most alarmingly, some grassland specialists, such as the enigmatic Great Indian Bustard and the Red-necked Falcon, experienced drastic reductions in their reporting rates (Figure 5).

#### Great Indian Bustard

Once widely distributed across Indian semi-arid grasslands, the Great Indian Bustard is now restricted to fragmented pockets of open habitats, with a steadily decreasing population (Dutta et al. 2011). Our results, consistent with previous studies from different parts of the country (Dutta et al., 2011; Narwade & Rahmani, 2020; Varghese et al., 2016) and national population trends, show that bustard presence in the region has declined during the study period (Figure 5). Their numbers have reached historic lows, and the species is nearing local extinction. Remote sensing and GIS studies have revealed that the suitable habitat for bustards in the Nannaj-Mardi region is extremely fragmented, with relatively small patches of grasslands remaining (Varghese et al. 2016).

Until 2016, the Great Indian Bustard showed distinct seasonality in the study area, being more frequently observed between June/July and October/November (Figure 5). Subsequently, its detection has been too low to discern seasonality. It is likely that the seasonal appearance of the bird at the study site reflects seasonality in the abundance of its prey: locusts, grasshoppers, beetles, frogs, bird eggs, small snakes and mice (Hume and Marshall 1879; Bhushan and Rahmani 1992; Patil et al. 2013), although this has not been examined specifically in our study area. SM notes that the breeding season for the bustard at Nannaj lasts from July/August to December each year, as evidenced by the presence of active nests and chicks during these months.

### B. Generalists

On the whole, generalist species showed increasing or indistinguishable from stable trends in their reporting rates (Figure 5). Most species showed trends indistinguishable from stable, albeit some with considerable fluctuation, over our 13-year study period. Generalists, by virtue of their ability to survive in varied environments, are likely to be less sensitive to changes in the landscape (Bowler et al. 2019; Callaghan et al. 2019).

A few species, such as the Short-eared Owl and Western Marsh Harrier (Figure 5), however, witnessed declines over time. These species, although generalists, are known to have a preference for more open habitats (Ali 1990), which are shrinking and increasingly fragmented at Nannaj (Varghese et al. 2016).

#### Indian Peafowl

The Indian Peafowl is a generalist species that has undergone considerable increases in our study period, consistent with trends from other places across the country (Figure 5). This is a species known to feed within agricultural lands and regularly cause crop loss (Paranjpe and Dange 2020), and is likely benefiting from recent agricultural expansions within the study region. However, peafowl increases at Nannaj are less dramatic than at the national level (SoIB 2020; Jose V and Nameer 2020).

### C. Human commensals

Our expectation was that increased urbanisation in recent years would result in an increase of all human commensal species over time, similar to their national trends (SoIB 2020). However, over the course of our whole study period, it is only the Feral Pigeon that shows considerable increases in its reporting rate. The other human commensal species (House Crow and Common Myna) show moderate long-term declines in the study region, while the House Sparrow is indistinguishable from stable.

#### Feral Pigeon

The Feral Pigeon (*Columba livia*) has undergone a dramatic increase in its abundance worldwide, closely correlated with increasing human density and urbanisation (Jokimäki and Suhonen 1998). Much of its recent increase can be attributed to the increase in human population and activity, the ability of the species to exploit diverse food sources in an urban setting, and reduced predation pressures within urban environments (Stukenholtz et al. 2019). Consistent with national and worldwide trends (Stukenholtz et al. 2019; SoIB 2020), our data shows rising reporting rates for Feral Pigeon within our study area (Figure 5).

## Conclusion

Our study uses a simple checklist method to understand changes in bird communities at a local scale. We reiterate that reporting rates are not synonymous with population densities. As shown by Altwegg and Nichols (2019), reporting rate tends to increase with population density but at a diminishing rate. This means that changes in reporting rate are likely to underestimate changes in population density, especially for abundant species. For this reason, trends in reporting rates must be viewed in light of the mean reporting rate of the species across the study period viz. how rare or common a species is.

Despite not estimating absolute population densities (which would require taking into account variation in detectability among species), checklist-based methods can provide important information on broad trends and provide early warnings of changes in populations, especially if one can assume constant within-species detectability over time. Of course, monitoring alone cannot diagnose causes of changes in abundance; but its results can trigger more detailed work towards that end.

The power of this method increases as the community of people in the practice of ecological monitoring grows. This includes birdwatchers (or other enthusiasts), Forest Department staff, nature guides, ecotourism outfits, college students, and many more. Daily bird lists (or lists of specific duration in a consistent location) can enable citizen scientists to understand their neighbourhood ecosystems (for example, the observations by Quader 2021) and Protected Area managers to monitor their lands. Online platforms, such as eBird, make it easy to record, store and visualise the collected data. Simple online tools for summarising and visualising the information collected would enable a more diverse set of people to take up this activity.

Our study demonstrates that even one dedicated individual, recording checklists consistently over a long time, can help our understanding of shifts in the local bird community. We look forward to a day when thousands of individuals across India keep track of the birds of their localities using simple, repeatable protocols such as that described here.

## Conflict of interest statement

The authors declare no conflict of interest.

## Acknowledgements

We are grateful to Ashwin Viswanathan for comments, criticisms and discussions throughout the writing process. Our thanks to Abinand Reddy for assistance and inputs during data cleaning. We thank the Maharashtra Forest Department for their encouragement and support. We would like to thank Sunil Limaye IFS, R N Nale, R N Kulkarni, B C Yele, K N Sabale, S K Jawale, Bharat Chedda, Shivakumar More, Aditya Kshirsagar, Pankaj Chindarkar, Balasaheb Lambture, Abhishek Deshpande, Sushant Kulkarni, and the staff of Range Forest Office (Great Indian Bustard Sanctuary).

## Financial support

Financial support for this work came from the Ministry of Environment, Forests and Climate Change, Government of India.

## Supplementary Material

**Figure S1:**
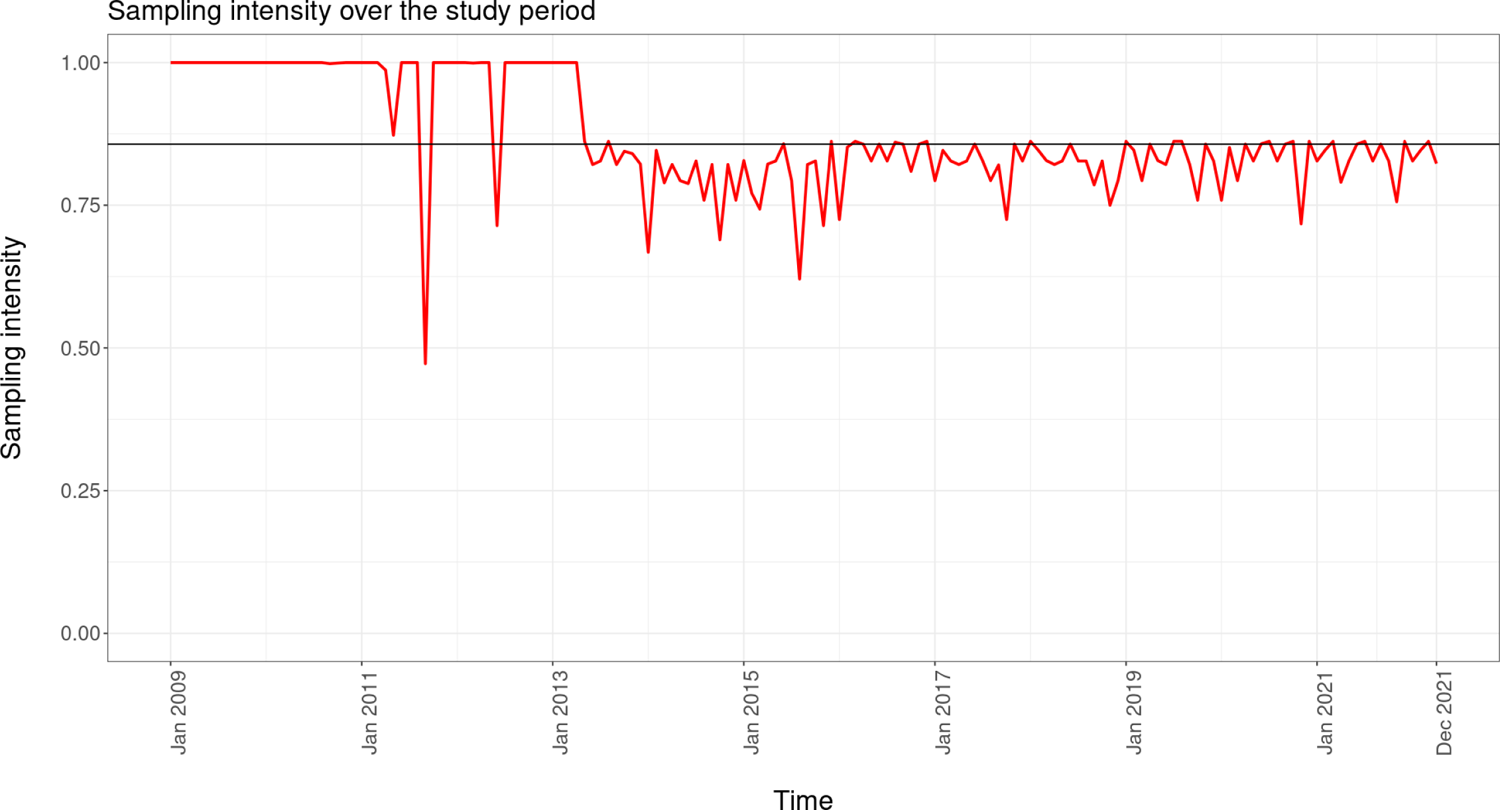
Sampling intensity of data over the study period, from 2009-2020. The sampling intensity was measured as the proportion of days of survey in each fortnight period. The horizontal line represents the mean sampling intensity throughout the study period.

## Notes

### Competing Interest Statement

The authors have declared no competing interest.

### Summary of Updates

We have revised our analysis and do not define any new indices of abundance, thus avoiding the complications arising from the propagation of SEs in aggregated quantities (as done earlier). Instead, we use a binomial GLMM to examine the trends of reporting rates over years, with months as the random effect. Further, we have also removed the normalisation of annual reporting rates with the first year. We have revised the plot showing species trends to highlight the mean reporting rates of the various species in our study. This revision will enable the reader to evaluate species trends in light of how commonly seen a species is.

## References

Ali S (1990) The book of Indian birds 11th ed. Oxford

Altwegg R, Nichols JD (2019) Occupancy models for citizen-science data. Methods Ecol Evol 10:8–21. https://doi.org/10.1111/2041-210X.13090

Bates D, Mächler M, Bolker B, Walker S (2015) Fitting Linear Mixed-Effects Models Using lme4. J Stat Softw 67:1–48. https://doi.org/10.18637/jss.v067.i01

Bhushan B, Rahmani AR (1992) Food and feeding behaviour of the great Indian bustard Ardeotis nigriceps (Vigors). J Bombay Nat Hist Soc 89:27–40

Billerman SM, Keeney BK, Rodewald PG, Schulenberg TS (eds) (2022) Birds of the World. Cornell Laboratory of Ornithology, Ithaca, NY, USA.

Bowler DE, Heldbjerg H, Fox AD, et al (2019) Long-term declines of European insectivorous bird populations and potential causes. Conserv Biol 33:1120–1130. https://doi.org/10.1111/cobi.13307

Brown LD, Cai TT, DasGupta A (2001) Interval Estimation for a Binomial Proportion. Stat Sci 16:101–117

Callaghan CT, Major RE, Wilshire JH, et al (2019) Generalists are the most urban-tolerant of birds: a phylogenetically controlled analysis of ecological and life history traits using a novel continuous measure of bird responses to urbanization. Oikos 128:845–858

Dutta S, Jhala Y (2014) Planning agriculture based on landuse responses of threatened semiarid grassland species in India. Biol Conserv 175:129–139. https://doi.org/10.1016/j.biocon.2014.04.026

Dutta S, Rahmani AR, Jhala YV (2011) Running out of time? The great Indian bustard Ardeotis nigriceps—status, viability, and conservation strategies. Eur J Wildl Res 57:615–625. https://doi.org/10.1007/s10344-010-0472-z

eBird (2021) eBird: An online database of bird distribution and abundance [web application]. eBird, Cornell Lab of Ornithology, Ithaca, New York. Available: http://www.ebird.org.

Fogarty DT, Elmore RD, Fuhlendorf SD, Loss SR (2017) Influence of olfactory and visual cover on nest site selection and nest success for grassland-nesting birds. Ecol Evol 7:6247– 6258. https://doi.org/10.1002/ece3.3195

Ganesh T, Prashanth M (2018) A first compilation of harrier roost counts from India suggests population declines of wintering birds over 30 years. Ardea 106:19–29

Gregory RD, van Strien A (2010) Wild bird indicators: using composite population trends of birds as measures of environmental health. Ornithol Sci 9:3–22

Grolemund G, Wickham H (2011) Dates and times made easy with lubridate. J Stat Softw 40:1– 25

Henwood WD (1998) An overview of protected areas in the temperate grasslands biome. Parks 8:3–8

Hume Allan Octavian, Marshall Charles Henry Tilson (1879) The game birds of India, Burmah, and Ceylon. Calcutta,A.O. Hume and Marshall

Jokimäki J, Suhonen J (1998) Distribution and habitat selection of wintering birds in urban environments. Landsc Urban Plan 39:253–263. https://doi.org/10.1016/S0169-2046(97)00089-3

Jose V S, Nameer PO (2020) The expanding distribution of the Indian Peafowl (Pavo cristatus) as an indicator of changing climate in Kerala, southern India: A modelling study using MaxEnt. Ecol Indic 110:105930. https://doi.org/10.1016/j.ecolind.2019.105930

Kher V, Dutta S (2021) Rangelands and crop fallows can supplement but not replace protected grasslands in sustaining Thar Desert’s avifauna during the dry season. J Arid Environ 195:104623. https://doi.org/10.1016/j.jaridenv.2021.104623

Kitowski I (2007) The roost and roosting behaviour of Eurasian Marsh Harriers Circus aeruginosus during autumnal migration in Eastern Poland. Pak J Biol Sci PJBS 10:734–740. https://doi.org/10.3923/pjbs.2007.734.740

Krishna YC, Kumar A, Isvaran K (2016) Wild Ungulate Decision-Making and the Role of Tiny Refuges in Human-Dominated Landscapes. PLOS ONE 11:e0151748. https://doi.org/10.1371/journal.pone.0151748

Madhusudan MD, Vanak A (2021) Mapping the distribution and extent of India’s semi-arid open natural ecosystems

Narwade SS, Rahmani AR (2020) Birds of the south-western Deccan Plateau region of Maharashtra, India, with special reference to the Great Indian Bustard Ardeotis nigriceps. 16:18

Paranjpe D, Dange P (2020) A tale of two species: human and peafowl interactions in human-dominated landscapes influence each other’s behaviour. Curr Sci 119:670

Patil P, Rahmani AR, Hallager S (2013) Behavioural ethogram of the Great Indian Bustard Ardeotis nigriceps (Vigors) 1831. J Bombay Nat Hist Soc JBNHS 110:22–34

Praveen J, Jayapal R, Pittie A (2016) A checklist of the birds of India. Indian BIRDS 11:113–170

Punjabi GA, Chellam R, Vanak AT (2013) Importance of Native Grassland Habitat for Den-Site Selection of Indian Foxes in a Fragmented Landscape. PLOS ONE 8:e76410. https://doi.org/10.1371/journal.pone.0076410

Quader S (2021) Neighbourhood bird monitoring through consistent listing. https://www.slideshare.net/suhelq/neighbourhood-bird-monitoring-through-consistent-listing. Accessed 3 Jun 2023

R. Development Core Team (2013) R: A language and environment for statistical computing Sethi VK, Bhatt D, Kumar A (2010) Hatching success in Yellow-wattled Lapwing Vanellus malabaricus. Indian Birds 5:139–142

SoIB (2020) First comprehensive assessment of bird species found in India. In: State Indias Birds. https://www.stateofindiasbirds.in/. Accessed 18 Jan 2022

Stukenholtz EE, Hailu TA, Childers S, et al (2019) Ecology of Feral Pigeons: Population Monitoring, Resource Selection, and Management Practices

Sullivan BL, Wood CL, Iliff MJ, et al (2009) eBird: A citizen-based bird observation network in the biological sciences. Biol Conserv 142:2282–2292. https://doi.org/10.1016/j.biocon.2009.05.006

Varghese AO, Sawarkar VB, Rao YLP, Joshi AK (2016) Habitat suitability assessment of Ardeotis nigriceps (Vigors) in great Indian bustard sanctuary, Maharashtra (India) using remote sensing and GIS. J Indian Soc Remote Sens 44:49–57

Verma A (1996) Harriers in India: a field guide. Wildl Inst India Dehradun India Organ Are Sincerely Thanked Assist Provid Field Stud Carried From

Wickham H (2016) Data analysis. In: ggplot2. Springer, pp 189–201

Wickham H, Averick M, Bryan J, et al (2019) Welcome to the Tidyverse. J Open Source Softw 4:1686

